# Hydroxychloroquine: mechanism of action inhibiting SARS-CoV2 entry

**DOI:** 10.1101/2020.08.13.250217

**Authors:** Zixuan Yuan, Mahmud Arif Pavel, Hao Wang, Scott B. Hansen

## Abstract

Hydroxychloroquine (HCQ) has been proposed in the treatment of SARS-coronavirus 2 (SARS-CoV-2) infection, albeit with much controversy. *In vitro*, HCQ effectively inhibits viral entry, but its use in the clinic has been hampered by conflicting results. A better understanding of HCQ’s mechanism of actions *in vitro* is needed to resolve these conflicts. Recently, anesthetics were shown to disrupt ordered monosialotetrahexosylganglioside1 (GM1) lipid rafts. These same lipid rafts recruit the SARS-CoV-2 surface receptor angiotensin converting enzyme 2 (ACE2) to an endocytic entry point, away from phosphatidylinositol 4,5 bisphosphate (PIP_2_) domains. Here we employed super resolution imaging of cultured mammalian cells to show HCQ directly perturbs GM1 lipid rafts and inhibits the ability of ACE2 receptor to associate with the endocytic pathway. HCQ also disrupts PIP_2_ domains and their ability to cluster and sequester ACE2. Similarly, the antibiotic erythromycin inhibits viral entry and both HCQ and erythromycin decrease the antimicrobial host defense peptide amyloid beta in cultured cells. We conclude HCQ is an anesthetic-like compound that disrupts GM1 lipid rafts similar to anesthetics. The disruption likely decreases viral clustering at both endocytic and putative PIP_2_ entry points.

**KEY POINTS:** *Question:* What is the molecular basis for antiviral activity of hydroxychloroquine?

*Findings:* Hydroxychloroquine disrupt lipid rafts similar to general anesthetics.

*Meaning:* Since lipids cluster ACE2 and facilitate viral entry, hydroxychloroquine appears to inhibit viral entry by disrupting the lipid clustering of the SARS-CoV2 receptor.

## INTRODUCTION

Coronavirus disease 2019 (COVID19), a viral infection caused by severe acute respiratory syndrome coronavirus 2 (SARS-CoV-2), recently emerged as a serious public health problem^1 2^. Currently, millions of people have been infected with SARS-CoV-2 worldwide. Proposed treatments for severe symptoms include a well-known FDA approved antimalarial agents chloroquine (CQ) and its derivative hydroxychloroquine (HCQ) ^3–7^, but their mechanism of action are poorly understood in COVID19 and their proven antiviral effects are largely limited to *in vitro* studies. A retrospective study claimed a benefit in particular with the macrolide antibiotic azithromycin^8^, but their use is not without controversy^9 10^, and randomized control studies without an antibiotic appeared to have no benefit^11^. In the treatment of malaria, CQ targets the replication cycle of the parasite^12^, a mechanism of action presumably unrelated to their action in COVID19. Understanding the underlying *in vitro* mechanism for these compounds in COVID19 could help in designing efficacious clinical trials and bettering the translation of their use in the clinic.

We have shown a cholesterol-dependent mechanism for anesthetics that regulates the movement of membrane proteins between monosialotetrahexosylganglioside1 (GM1) containing lipid rafts and PIP_2_ lipid domains^13 14^. The GM1 rafts are formed by cholesterol packing^15^ and the PIP_2_ domains are formed from charged protein clustering^16^ (Fig. S1A). In cellular membranes, local and general anesthetics, including propofol, disrupt GM1 rafts^13 17^. The anesthetic propofol also has various beneficial effects on COVID19 treatment and the FDA recently permitted the emergency use of Fresenius Propoven 2% emulsion to maintain sedation via continuous infusion for COVID-19 patients^18^.

Cholesterol is critical to both viral entry and immune responses^19^. We recently showed the SARS-CoV-2 surface receptor, angiotensinogen converting enzyme 2 (ACE2)^20–22^ moves between GM1 rafts and PIP_2_ domains in a cholesterol dependent manner^23^. In an obese mouse model, cholesterol was high in lung tissue and this correlated with ACE2 movement to endocytic lipids, a condition that accelerated viral entry into the target cells in cell culture^23^.

Interestingly, CQ is an anesthetic—subcutaneous injection of CQ produces instant local anesthesia sufficient to perform a surgical procedure^24 25^ and structurally CQ is strikingly similar to a local anesthetic (Fig. 1A). Both CQ and local anesthetics such as tetracaine are weak bases and their uptake changes the acid base balance within the membrane^26 27^. Additionally, common local anesthetics such as mepivacaine, bupivacaine, tetracaine and other lipid raft disrupting compounds, such as sterols and cyclodextrin, can exert anti-viral or anti-microbial activity^28–31^.

**Fig. 1.**
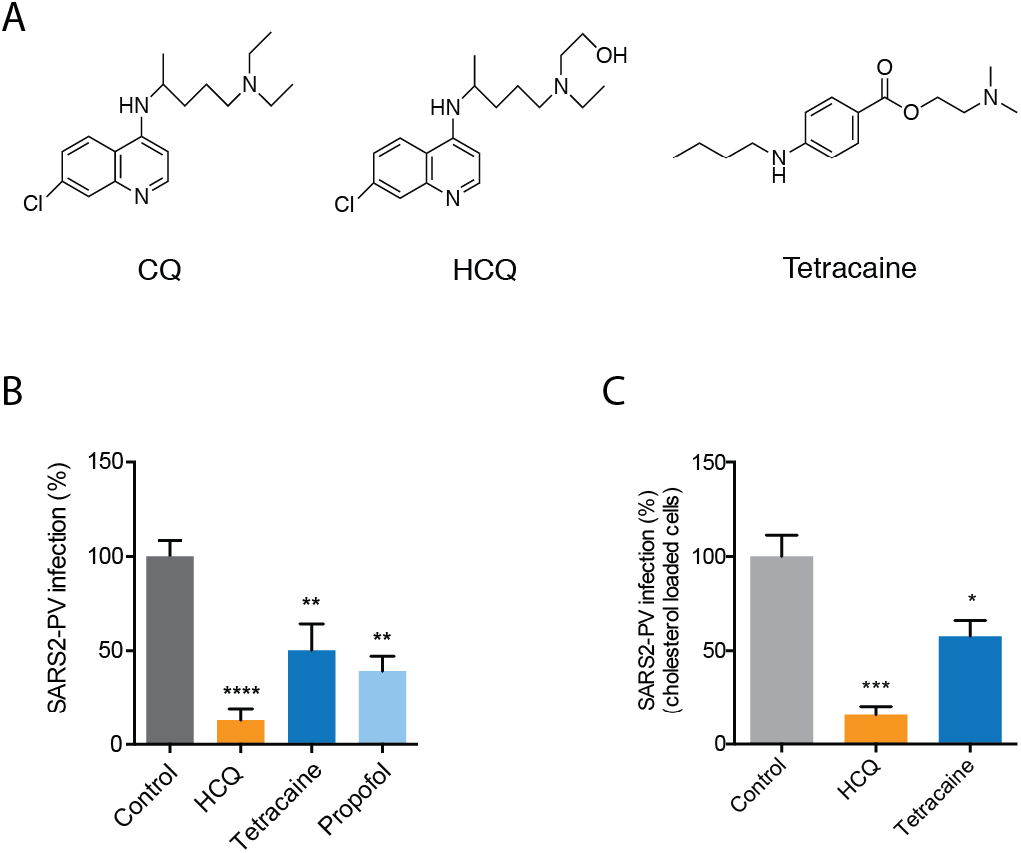
Anesthetics and hydroxychloroquine inhibit SARS2-PV entry. **(A)** Chemical structures comparing chloroquine (CQ) and hydroxychloroquine (HCQ) to tetracaine, a local anesthetic. **(B-C)** SARS-CoV-2 pseudovirus (SARS2-PV) entry assay measured as a percent of control luciferase activity. HCQ (50 μM), tetracaine (50 μM) and propofol (50 μM) inhibited viral infection in ACE2 overexpressing HEK293T cells without (B) and with (C) cholesterol loading (4 μM apolipoprotein E + 10% serum). Data are expressed as mean ± s.e.m., *P ≤ 0.05, **P ≤ 0.01, ***P ≤ 0.001, ****P ≤ 0.0001, one-way ANOVA, n=4-6.

These properties led us to compare the drugs’ lipid disruption properties with viral entry and potentially understand a component of their underlying molecular mechanism. Understanding CQ’s mechanism of action could help understanding potential contradictions and determining which cellular and animal models are appropriate for testing CQ’s effect *in vitro* and *in vivo*.

Here we employed super resolution imaging to show that HCQ disrupts GM1 rafts in a manner similar to anesthetics causing ACE2 to leave GM1 rafts (the endocytic site of viral entry) and PIP_2_ domains.

## RESULTS

### Inhibition of SARS-CoV2 entry by anesthetic-like compounds

In order to test SARS-CoV2 viral entry, we expressed a retrovirus pseudotyped with the SARS2 spike protein (SARS2-PV) in HEK 293T cells. A segment of the spike protein binds to ACE2 and recapitulates viral entry^32 33^. A luciferase encoded in the pseudotyped virus allows quantitative measurement of viral entry.

Treatments with propofol, tetracaine, and HCQ all robustly reduced SARS2-PV entry into HEK293T cells overexpressing ACE2 receptor (Fig. 1B). The cells were first treated with drug (50 μM) and then the drug was removed, after which SARS2-PV was applied (*i.e.*, the virus did not experience the drug directly, only the cells). HCQ had the greatest effect with almost a 90% reduction in SARS2-PV luciferase activity (Fig. 1B).

Since COVID19 is often severe in obese patients, we also tested HCQ inhibition in cells loaded with cholesterol. To load cells with cholesterol, we treated HEK293T cells overexpressing ACE2 receptor with 4 μM apolipoprotein E (ApoE) and 10% fetal bovine serum. ApoE is a cholesterol carrier protein linked to the severity of COVID19^34^. In tissue, ApoE binds to low-density lipoprotein (LDL) receptor and facilitates loading of cholesterol into cells^35^ (Fig. S1B). When apoE is in excess in low cholesterol conditions, it facilitates efflux of cholesterol from the cell^35^. To provide a source of cholesterol to the apoE, we added 10% fetal bovine serum (FBS), a common source of cholesterol ~310 μg/mL. Importantly, apoE is not present in FBS^36^ allowing us to carefully control cholesterol loading. Cells were treated acutely (1 hr.) prior to viral infection. We adapted cholesterol loading from our previous studies in primary neurons ^37^.

Loading cells with cholesterol into HEK293T cells overexpressing ACE2 receptor increased viral entry by 56% (Figure S1C), consistent with our previous studies where cholesterol loading increased viral entry by ~50% (p value <0.05) with endogenously expressed ACE2^23^. Control cells in ACE2 overexpressed cell line were highly variable in viral infection, reducing statistical significance. Nonetheless, as expected, treatment of cholesterol loaded cells with HCQ and tetracaine reduced SARS2-PV entry in high cholesterol. The efficacy with cholesterol was slightly less compared to non-cholesterol loaded cells.

### HCQ acts in an anesthetic pathway by disrupting lipid rafts

Based on the structural similarities of HCQ with anesthetics (Fig. 1A), we investigated HCQ’s ability to work through an anesthetic-like mechanism. We recently showed a mechanism of anesthesia based on disruption of lipid rafts. Anesthetics perturb rafts in two ways. First, anesthetics increase the apparent size and number of lipid rafts as observed using super resolution imaging and cluster analysis^13 17^. Second, anesthetics can disassociate cholesterol sensitive proteins from GM1 rafts. The disassociation of a proteins from a GM1 raft can also be measured with super resolution imaging using two-color fluorescent labeling and pair correlation analysis. The results of pair correlation are quantitative, well established, and provide mechanistic insights into raft associated signaling and protein movement at nanoscale distances^13 17^.

To test HCQ’s effects on lipid membranes, we first examined the effect of HCQ on the apparent structure (size and number) of GM1 rafts by direct stochastic optical reconstruction microscopy (dSTORM) in the membranes of HEK293T cells using density-based spatial clustering of applications with noise (DBSCAN). We found 50 μM HCQ, a minimum saturating concentration that was shown to inhibit viral entry^3^, increased the number and apparent size (Fig. 2A-C) of GM1 rafts. HCQ’s perturbation was very similar to that previously seen for general anesthetics chloroform and isofluorane^13^ (Fig. 2A). We also observed Ripley’s H(r) of GM1 clusters decreased after HCQ treatment (Fig. S2A), suggesting a decrease in domain separation. Methyl-beta cyclodextrin (MβCD), a chemical known to deplete GM1 rafts from the cell membrane, decreased both the number and apparent size of GM1 clusters (Fig. 2A-B).

**Fig. 2.**
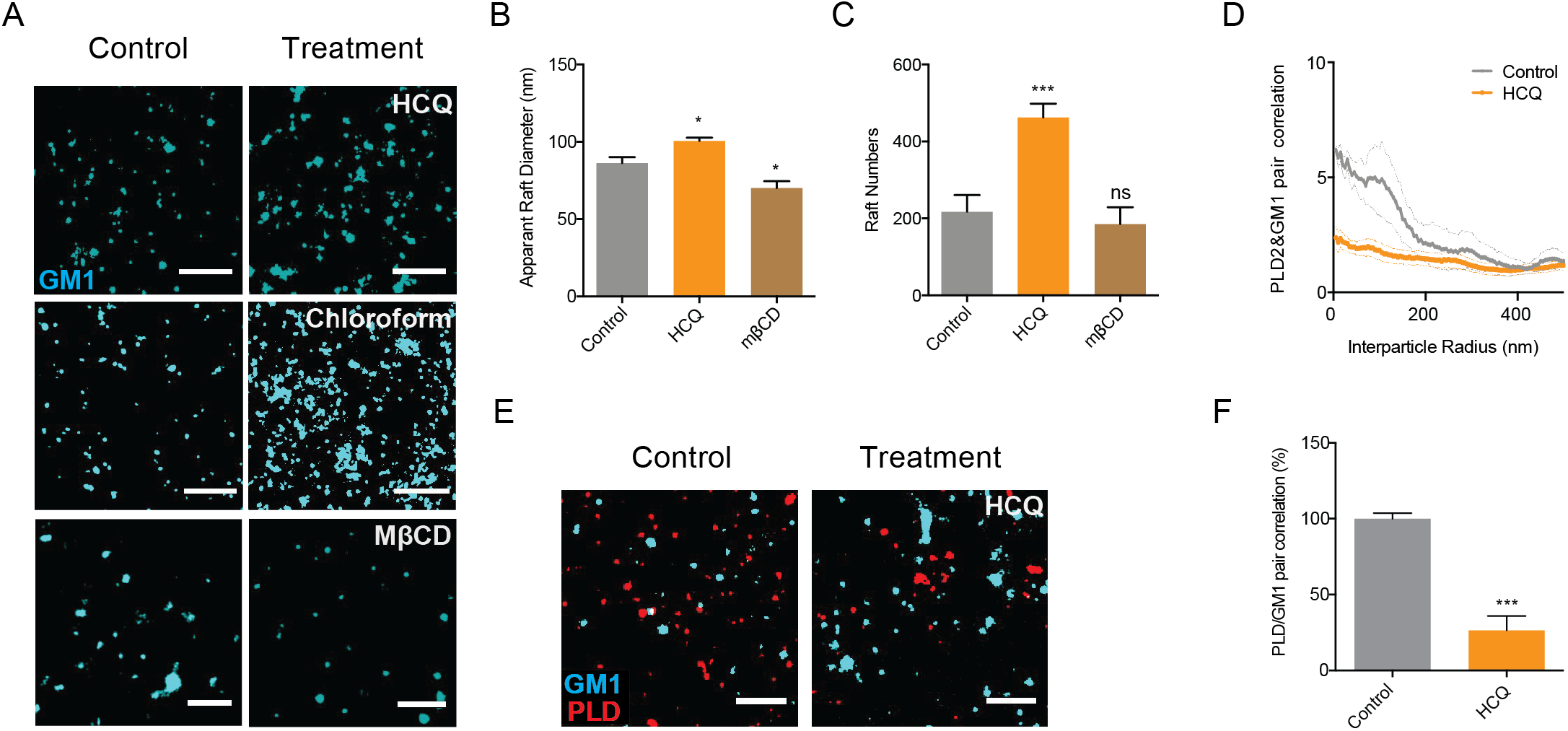
Anesthetic-like disruption of GM1 rafts by Hydroxychloroquine. **(A)** Representative dSTORM images showing the GM1 raft perturbation by HCQ (50 μM) and MβCD (100 μM) in HEK293T cells (Scale bars = 1 μm). Similar disruption from 1 mM chloroform treatment, an anesthetic, is shown from Pavel et. al. PNAS 2020; 117:13757–66, with permission, for comparison. **(B-C)** Bar graph of the apparent raft diameter analyzed by DBSCAN cluster analysis. HCQ increases both raft diameter (B) and number (C) of GM1 rafts. Data are expressed as mean ± s.e.m., *P ≤ 0.05, ***P ≤ 0.001, one-way ANOVA, n=7. **(D-E)** Pair correlation analysis (D) of two color dSTORM imaging (E). HCQ treatment decreased association of phospholipase D2 (PLD2, red shading), an anesthetic sensitive enzyme, with GM1 rafts (cyan shading) (scale bars = 1 μm). **(F)** Quantification of pair correlation in (D) at short distances (0-5 nm). Data are expressed as mean ± s.e.m., ***P ≤ 0.001, unpaired t-test, n=4-7.

To test HCQ for anesthetic-like properties, we treated HEK293T cells with 50 μM HCQ, labeled GM1 lipids and the protein phospholipase D2 (PLD2) with (CTxB, a pentadentate toxin that labels GM1 lipids and anti PLD2 antibody respectively). Phospholipase D2 (PLD2) is an anesthetic sensitive enzyme that translocates out of GM1 rafts in response to general anesthetics xenon, chloroform, isoflurane, propofol, and diethyl ether and provides a live cell test for disruption of lipid rafts^13 38^.

We found 50 μM HCQ robustly disrupted PLD2 localization with GM1 rafts (Fig. 2D-E). Quantification of the % pair correlation at short radiuses (0-5 nm) decreased by almost 70% (Fig. 2F). Hence HCQ’s effect on the lipid membrane is clearly similar to general anesthetics (Fig. 2A) where the domains are larger but the ability to retain a palmitoylated protein is inhibited. We attempted to confirm HCQ disruption of lipid rafts in a live-cell PLD2 activity, however HCQ appeared to directly inhibit enzymatic activity (Fig. S2B-C) precluding the assays usefulness. Nonetheless, HCQ’s displacement of PLD2 observed by super resolution imaging led us to consider displacement of the ACE2 receptor.

### HCQ disrupts clustering of ACE2 with GM1 rafts

The ability of a virus to cluster is important for its infectivity and maturation^39–42^. For SARS-CoV2 entry depends on binding to ACE2. ACE2 localizes to both GM1 rafts and PIP_2_ domains, however in high cholesterol (obese animals) ACE2 is in GM1 rafts^23^. To recapitulate a physiologically relevant environment of an obese patients, we loaded HEK293T cells with cholesterol using apoE and serum identical to the treatment in Figure 1^23^. The cells were then fixed, labeled them with anti-ACE2 antibody and CTxB, and imaged using two color dSTORM.

After 50 μM HCQ treatment, we found ACE2 receptor dramatically decreased its association with GM1 rafts, despite the increase in GM1 raft size and number (Fig 3A). Pair correlation was decreased at all distances (Fig. S3A). At short distances (0-5 nm) the decreased was almost 50% (p<0.05) (Fig. 3B) confirming HCQ disrupts the ability of GM1 rafts to sequester ACE2 as expected from its anesthetic-like mechanism of action and its effect on PLD2.

**Fig. 3.**
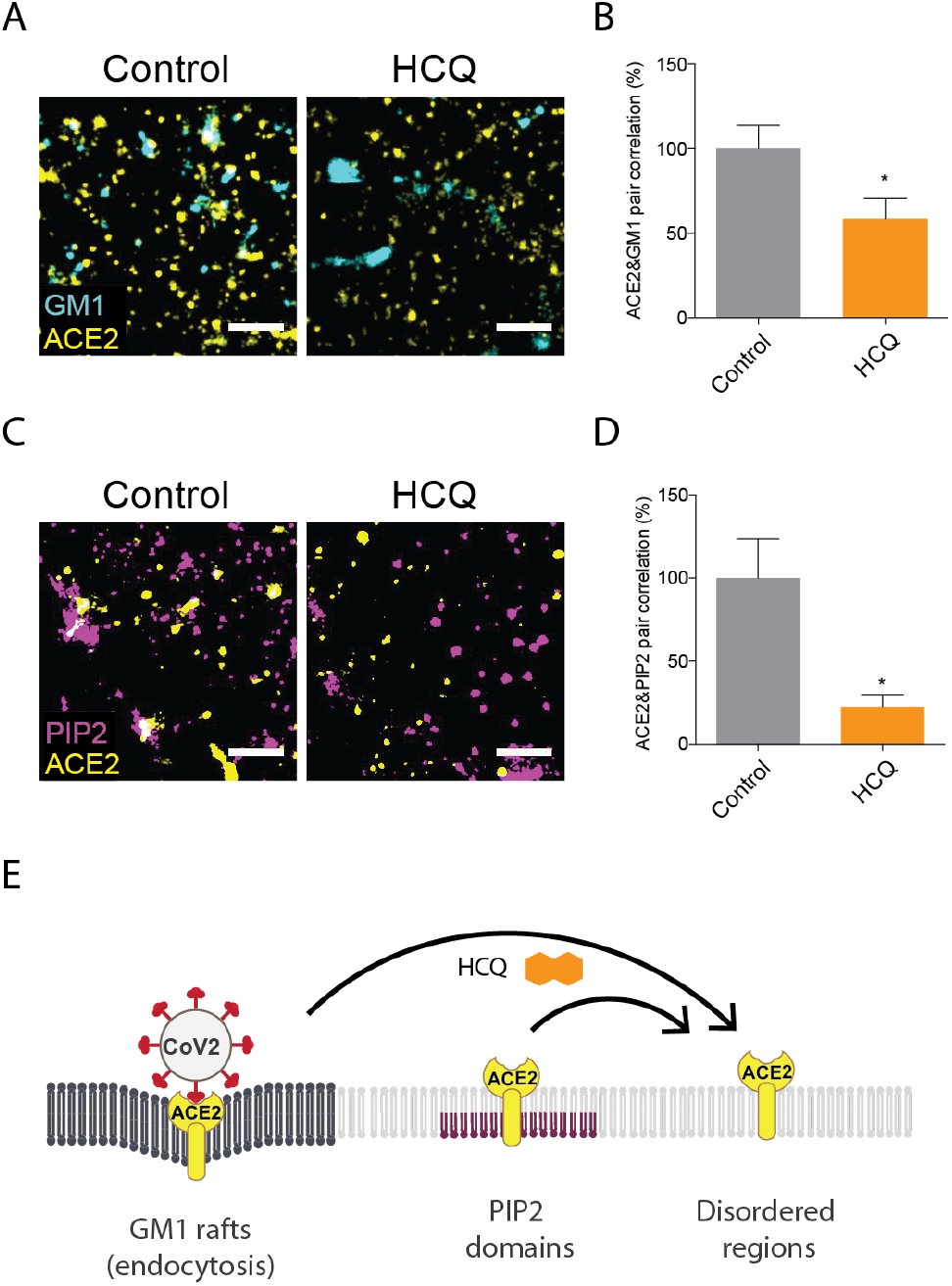
Hydroxychloroquine moves ACE2 from GM1 rafts and PIP_2_ domains. **(A)** Representative dSTORM super resolution images showing the effect of HCQ (50 μM) on the nanoscale localization of ACE2 (yellow) with GM1 rafts (cyan) after loading HEK293T cells with cholesterol (scale bars = 1 μm). **(B)** Percent of pair correlation (Fig. S3A) calculated at short distances (0-5 nm). HCQ decreased the pair correlation between ACE2 and GM1 rafts indicating a decrease in association between PLD and GM1 rafts. Data are expressed as mean ± s.e.m., *P ≤ 0.05, unpaired t-test, n=6. **(C)** Representative dSTORM super resolution images of ACE2 (yellow) and PIP_2_ domain (magenta) in HEK293T cells at normal cholesterol level after the treatment of HCQ (50 μM) (scale bars = 1 μm). **(D)** HCQ decreased the pair correlation between ACE2 and PIP_2_ domains indicating a decrease in association between PLD and PIP_2_ domains. Data are expressed as mean ± s.e.m., *P ≤ 0.05, unpaired t-test, n=5. **(E)** Model showing HCQ (orange hexagon) inducing translocation of ACE2 (yellow receptor) from GM1 rafts (dark grey lipids) in high cholesterol. HCQ disrupts ACE2 interaction with PIP_2_ domains causing ACE2 to translocate to the disordered region.

### HCQ disrupts PIP_2_ domains

We have previously shown that disrupting GM1 rafts moves ACE2 to and clusters with PIP_2_ domains^23^. We presume PIP_2_ domains reside in the disordered regions away from GM1 rafts due to the large amount of unsaturation in PIP_2_’s acyl chains^43 44^. To determine whether ACE2 clusters with PIP_2_ domains after disruption of GM1 rafts, we co-labeled ACE2 and PIP_2_ domains in HEK293T cells at normal cholesterol levels and treated the cells with and without 50 μM HCQ. Figure 2D shows representative figures of dSTORM imaging. Somewhat unexpected, the pair correlation between ACE2 and PIP_2_ domains clearly decreased with HCQ treatment (Fig. 2E), suggesting HCQ disrupts ACE2 association with both GM1 rafts and PIP_2_ domains.

We further characterized the nature of the PIP_2_ disruption by analyzing the structures of the PIP_2_ domains before and after HCQ treatment using dSTORM cluster analysis. Figure 2D shows representative dSTORM images comparing PIP_2_ domains before and after HCQ treatment. HCQ treatment decreased both the number and size of PIP_2_ domains by ~ 25% and 50% respectively (Fig. S3B-C) confirming HCQ directly perturbs the domains.

### Erythromycin inhibits viral entry through perturbing GM1 rafts

Azithromycin, an antibiotic derived from erythromycin, has been given to COVID-19 patients in combination with HCQ and was shown to significantly reduce COVID-19 associated mortality^8^. Erythromycin has shown antiviral properties in numerous studies^45–48^. Many antimicrobials are also thought to disrupt GM1 rafts. Based on the cholesterol sensitivity of SARS-CoV2 we reasoned erythromycin could contribute to an antiviral effect through raft disruption leading us to test its effect on SARS2-PV.

We found erythromycin (100 μg/mL) inhibited SARS2-PV infection ~70% in HEK293T cells over expressing ACE2 at normal cholesterol levels (Fig. 3A). Consistent with disruption, the same treatment increased membrane fluidity ~ 60% (Fig. 4B). Furthermore, using PLD2 activity as a surrogate for an effect in live cells showed an ~10% increase (Fig. 4C), consistent with raft disruption. Unexpectedly, when we increased the cholesterol level using apoE and serum, erythromycin was no longer effective. In fact, when we examined the pair correlation of ACE2 and GM1 correlation we saw increased association of ACE2 with GM1 (Fig. S3D-E). This suggests the effect of erythromycin on viral entry may be bimodal and depend on the progression of the disease. If HCQ reverses the effects of cholesterol on disruption this may explain some of the proposed combined benefit of HCQ with erythromycin.

**Fig. 4.**
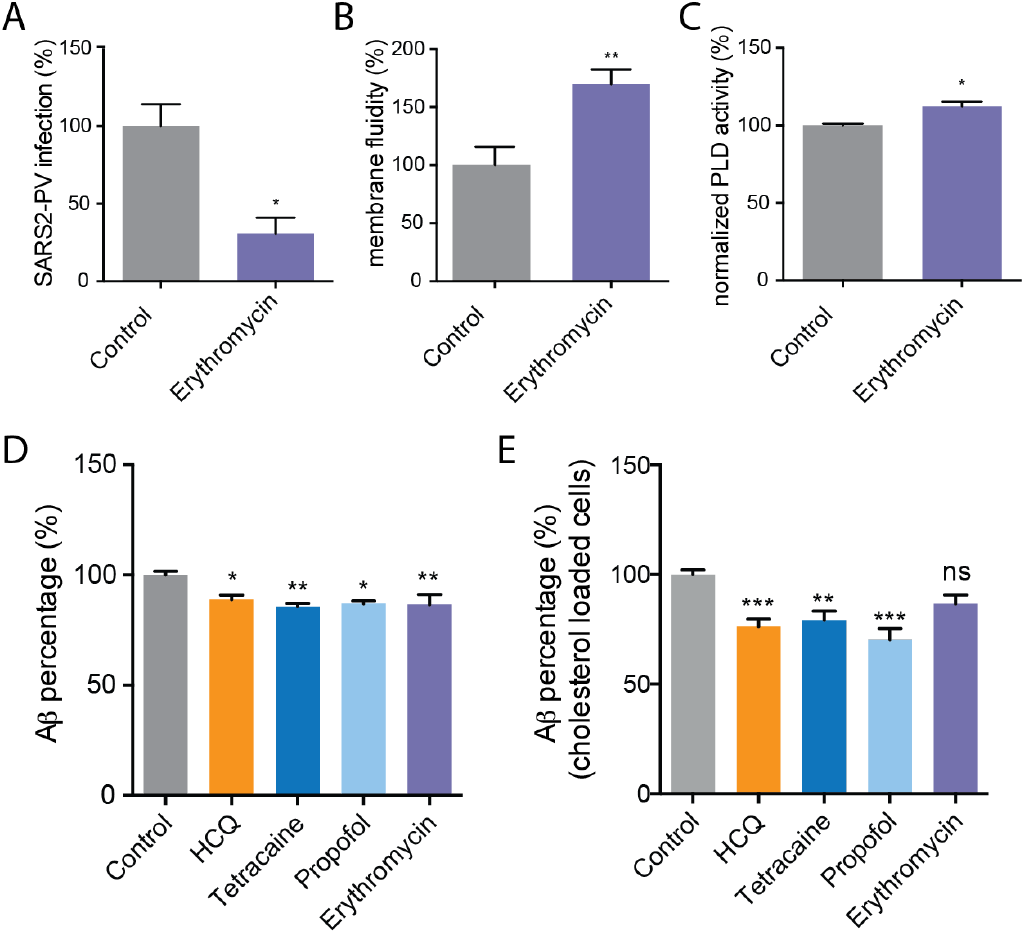
Erythromycin inhibits SARS-CoV-2 viral entry. **(A)** Percent SARS-CoV-2 pseudovirus (SARS2-PV) infection after erythromycin (100 μg/ml) treatment of HEK293T cells over expressing ACE2. Data are expressed as mean ± s.e.m., *P ≤ 0.05, unpaired t test, n=3. **(B)** Erythromycin (100 μg/ml) increased membrane fluidity in membrane fluidity assay. Data are expressed as mean ± s.e.m., **P ≤ 0.01, unpaired t test, n=3-4. **(C)** A raft disruption assay based on PLD2 enzymatic activity in HEK293T cells. **(D-E)** An ELISA assay showing HCQ (50 μM), tetracaine (50 μM), propofol (100 μM), and erythromycin (100 μg/ml) decreased the synthesis of Aβ40 in HEK293T cells with (E) and without (D) cholesterol loading (4 μM apolipoprotein E + 10% serum). Data are expressed as mean ± s.e.m., *P ≤ 0.05, **P ≤ 0.01, one-way ANOVA, n=3-7.

### HCQ’s disruption on host defense peptides

Lastly, we considered HCQ effect on host defense peptides. Host defense peptides are amphipathic antimicrobial peptides that are upregulated during an immune response and perturb the membranes of microbes^49 50^. Cholesterol and raft integrity are of great importance to the modulation of both innate and acquired immune responses^51^. Amyloid-beta (Aβ) has been demonstrated to protect against microbial infection as a host defense peptide and the production of Aβ is regulated by lipid raft integrity^37 52^ (Fig. S1D). We hypothesized that if HCQ disrupts lipids, it may disrupt the production of host defense peptides. Disrupting host defense peptides would be an unwanted effect that would need to be considered when treating COVID19 patients.

To investigate the role of HCQ and anesthetics-induced raft perturbation in the synthesis of host defense peptides, we measured Aβ production using a sandwich enzyme-linked immunosorbent assay (ELISA). We found that HCQ reduced Aβ generation ~ 10% in cultured HEK293T cells (Fig. 4D). The effect was statistically significant (p<0.05). We then loaded HEK293T cells with cholesterol (apoE + serum) to better reflect the disease state of COVID-19 with severe symptoms. The reduction of Aβ was ~20% greater in high cholesterol. Since tetracaine and propofol also disrupt GM1 rafts, we tested their effect on Aβ production and found it to be very similar in both high and low cholesterol. Interestingly, erythromycin did disrupt Aβ production in low cholesterol, but in high cholesterol the effect was very small and not statistically significant.

## Discussion

Taken together our finding shows a component of HCQ acts through anesthetic-like mechanism that disrupts ACE2 localization at both GM1 rafts and PIP_2_ domains. Presumably the disruption of ACE2 clustering decreases the ability of the virus to cluster and enter the cell. Figure 5 shows a propose a model of HCQ disrupting SARS-CoV-2 viral entry through raft perturbation. In an inflamed state, cholesterol traffics ACE2 from PIP_2_ domains to GM1 rafts where virions dock and enter through endocytic pathway. The perturbation of both GM1 rafts and PIP_2_ domains by HCQ likely inhibits viral entry by making it more difficult to cluster ACE2 and enter the endocytic entry point. The mechanism of surface receptor clustering was recently shown to be important to the related influenza virus^39^. How the molecular studies related to the clinic will need to be studied.

**Fig. 5.**
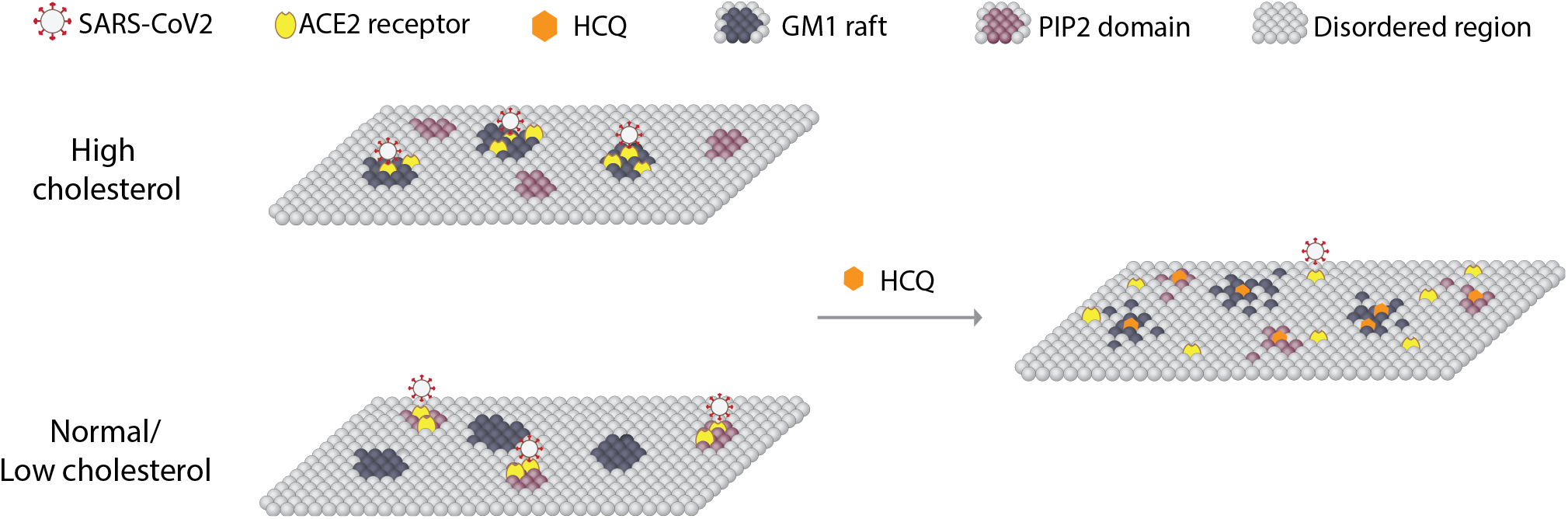
Model for HCQ mechanism of action in SARS-CoV-2 infectivity. A representation of the plasma membrane shows nanoscale lipid heterogeneity. Saturated monosialotetrahexosylganglioside1 (GM1) lipid rafts (dark grey) attract ACE2 (yellow oval) in high cholesterol (top). In low or normal cholesterol, ACE2 associates primarily with phosphatidylinositol 4,5-bisphosphate (PIP_2_). Hydroxychloroquine (HCQ, orange hexagon) disrupts the lipid order and excludes the association of ACE2 from both GM1 rafts and PIP_2_ domains (right panel). The SARS-CoV-2 virus (white circle with red spike) binds to the ACE2 receptor. The location of the ACE2 receptor dictates the location and efficacy of viral entry.

In part, HCQ counteracts cholesterol. The immune system uses cholesterol as a signal to fight infection^51^, including the release of peptides that kill bacteria (e.g., host defense peptides like Aβ)^49 50^. Not surprisingly, adding drugs that counteract the cholesterol to reduce viral entry reduce Aβ. The results of HCQ and anesthetics reducing the production of Aβ provide a molecular rational to supply exogenous antibiotics in combination with HCQ so that the overall antibiotics level is maintained.

To date only one large randomized controlled study has included azithromycin. Overall improvement to clinical scores was not statistically significant, but death with both HCQ and azithromycin were decreased (3 compared to 5 for control)^53^. Randomized control trials with HCQ alone have shown no benefit^11^. It is not clear at this time why *in vitro* and *in vivo* experiments with HCQ differ. Anesthetics often require mechanical ventilation, suggesting studies should focus on death outcome not mechanical ventilation. None of the studies considered lung cholesterol. If HCQ acts through cholesterol, then stratifying patients by lung cholesterol level may reveal a benefit. Cholesterol is typically measured in the blood not the tissue. The blood is primarily a transient transport system and may not accurately predict years of accumulation in the lung of obese or chronically inflamed patients^23 51^.

When cholesterol is low, HCQ likely has a reduced effect, i.e., the drug likely fails to reverse high cholesterol in the absence of high cholesterol. Hence, animal and cultured cell experiments in low cholesterol likely fail to capture the full benefit of HCQ and should be carefully scrutinized. For example, lungs cell and monkey’s showed no reduced effect from HCQ were performed without high cholesterol, but this reflects the physiological state of a child or healthy adult, does not reflect an obese patient at risk for death from severe COVID19 symptoms^54 55^. Over expressing a protease that primes the virus appears to overcome the antiviral properties of HCQ^54^. Our data suggests this is due to viral priming in the disordered region due to very high concentrations of the protease. Further studies are need to determine if the virus can be cleaved and enter a cell while in the disordered region with physiological levels of protease.

HCQ anesthetic-like properties are likely enhanced by its positively charged amine. HCQ appears to interact directly with PIP_2_ to block ACE2 localization with PIP_2_. It is unclear where ACE2 resides when it is excluded from both GM1 rafts and PIP_2_ domains. Presumably it moves into a generic disordered region of the cell membrane. Alternatively, it may move into PIP_3_ domains. PIP_3_ is typically short chain saturated^43 44^ and could possibly attract ACE2 if HCQ preferentially disrupts long chain polyunsaturated lipids such as PIP_2_.

Erythromycin, an analog of azithromycin, also contains a tertiary amine. Other aminoglycosides (e.g. neomycin) are known to bind tightly to and scavenge PIP_2_. Scavenging PIP_2_ is normally thought to block ligand binding^56^ or change a surface charge. Our data here suggests hydrophobic charged molecules disrupt PIP_2_ and the resulting ACE2 clustering. We previously assumed the inhibition of PLD2 by tetracaine was through direct binding of tetracaine to the enzyme, but here HCQ did not inhibit purified cabbage PLD (Fig. S2D), suggesting the inhibition could also occur through disruption of PIP_2_ and its ability to bind PLD2.

A previous mechanism suggested that HCQ could inhibit SARS-COV-2 viral entry step by changing the glycosylation of membrane proteins^57 58^. Our raft-associated protein activation mechanism is consistent with changes in glycosylation if the glycosylated protein is also sensitive to localization in lipid rafts. Many proteins are regulated by palmitoylation and PIP_2_, including numerous inflammatory proteins.

Based on the significant inhibition of SARS2-PV entry from tetracaine and propofol, a local anesthetic and a general anesthetic, anesthetic-like chemicals are likely helpful to treating COVID-19. Anesthetic with sugammadex dramatically decreases post-operative pulmonary complication^59^. The similarities between HCQ and anesthetics in chemical structure, viral entry inhibition, and raft perturbation tempt us to hypothesize that both HCQ and anesthetics share a parallel mechanism of action. Consistent with this hypothesis CQ has side effects similar to those reported in anesthetics^60^.

The increased viral infection in ACE2 over expressed cells is likely due to increased number of total lipid rafts not a shift from ACE2 into GM1 domains. Over expression of ACE2 causes loss of raft regulation and the enzyme likely distributes into both raft and no-raft regions^23^, suggesting over expression is a completely unphysiological condition. Hence the reduced efficacy of tetracaine and HCQ inhibiting viral entry in high cholesterol (57.7% vs 50.3% and 15.6% vs. 12.8% respectively, see Fig. 1C) cannot be directly attributed to a physiologically relevant mechanism. Nonetheless, in wild type cells ACE2 localization is still sensitive to cholesterol and the same trend should hold true. This conclusion is further supported in Fig. 4d-e where erythromycin was unable block AB production in high cholesterol.

All the imaging was performed with endogenously expressed proteins to ovoid loss of raft associated regulation of ACE2. The lipids were labeled after fixing to reduce movement between domains during labeling and to limit potential local lipid clustering by CTxB, especially saturated lipids. CTxB is pentadentate and in unfixed lipids causes clustering^61^ and to some degree CTxB clustering occurs in fixed cells^62^. Since we examined disruption of lipids, the CTxB could have decreased the amount of disruption we reported for apparent GM1 raft size (i.e., the amount of disruption may be under reported). PIP_2_ is polyunsaturated and we expect it is much better fixed in the membrane.

## METHODS

### Reagents

Hydroxychloroquine was purchase from Cayman Chemical and tetracaine was purchased from Sigma-Aldrich. Purified PLD2 from cabbage was purchased from Sigma-Aldrich respectively. PLD assay reagent amplex red 10-Acetyl-3,7-dihydroxyphenoxazine and 2-dioctanoyl-sn-glycero-3-phosphocholine (C8-PC) were purchased from Cayman Chemical. Horseradish peroxidase and choline oxidase were purchased from VWR. Methylbetacyclodextrin (MβCD) was purchased from Sigma-Aldrich.

### Psuedo-typed SARS-CoV-2 (SARS2-PV) Viral Entry Assay

#### Cells and virus

HEK293T cell line was cultured in Dulbecco’s Modified Eagle Medium (DMEM) with 10% fetal bovine serum (FBS) at 37 °C with 5% CO_2_ atmosphere. SARS-CoV-2 pseudotyped particles were constructed using plasmid co-transfection, and the particles were maintained at −80°C. The constructs were a gift from Dr. Mike Farzan, Scripps Research, Florida. Evaluation of antiviral activities HEK293T ACE2 overexpression cells (0.5 × 105 cells/well), also provided by Dr. Mike Farzan, were cultured in 96-well cell-culture plates (Corning™ Coastar™ Cell Culture 96 well plates, #3585) were incubated with 100 μL pseudotyped particles of each type, together with 50 μM hydroxychloroquine sulfate (HCQ, Cayman, #17911) or 50 μM tetracaine hydrochloride (Sigma-Aldrich, #T7508) for 1 h. Then, the virus-drug mixture was removed, and fresh medium was added. After 24 h, the particles yields were determined through a luciferase assay. Cells were washed with PBS and 16 μL Cell Culture Lysis Reagent (Promega, #E153A) was added into each well. The plate was incubated for 15 min with rocking at room temperature. 8 μL of cell lysate from each well was added into a 384-well plate, followed by the addition of 16 μL of Luciferase Assay Substrate (Promega, #E151A). Luciferase activity measurement was performed on a Spark 20M multimode microplate reader (Tecan). The luciferase activity as infection yields were plotted in GraphPad Prism 6 software. All the infection experiments were performed in a biosafety level-2 (BLS-2) laboratory.

### Super Resolution Microscopy (dSTORM)

To detect the lipid raft perturbation by hydroxychloroquine we employed Super Resolution Microscopy as described previously^14^. Briefly, HEK293T cells were grown in 8-well chamber slides (Nunc Lab-Tek chamber slide system, Thermo Scientific), washed and treated with 30-50 μM hydroxychloroquine for 30 min. The cells were then fixed with 3% paraformaldehyde, 0.1% glutaraldehyde, 30-50 μM hydroxychloroquine for 20 min, quenched with 0.1% NaBH_4_ for 7 min. Cells were then washed with PBS (three times) and permeabilized with 0.2% Triton-X 100 in PBS for 15 min. The permeabilized cells were blocked using a standard blocking buffer containing 10% BSA and 0.05% Triton in PBS for 90 min. For labelling, cells were incubated with primary antibody (anti-ACE2 antibody (Abcam, #ab189168), anti-PLD2 antibody, or anti-PIP_2_ antibody) for 60 min in antibody buffer (PBS with 5% BSA and 0.05% TritonX-100) at room temperature followed by 5 washes with wash buffer (PBS with 1% BSA and 0.05% TritonX-100) for 15 min each. Secondary antibodies (donkey anti-rabbit Cy3B and Alexa 647 conjugated CTxB) were added with antibody buffer for 30 min at room temperature followed by 5 washes as stated above. Then, cells were washed with PBS for 5 min and fixed for 10 min with fixation buffer as above, followed by 5 min washes with PBS for 3 times and 3 min washes with deionized distilled water. All steps except for pre- and post-fixation were performed with shaking.

A Zeiss Elyra PS1 microscope was used for super resolution microscopy with an oil-immersed 63X objective lens in TIRF mode. Images were acquired by Andor iXon 897 EMCCD camera and Zen 10D software with an exposure time of 18 ms per acquisition. Total 7,000-10,000 frames were collected. Alexa Fluor 647 and Cy3B were excited with a 642 nm and 561 nm laser in a photo-switching buffer consisting 1% betamercaptoethanol, 0.4 mg glucose oxidase and 23.8 μg catalase (oxygen scavengers), 50 mM Tris, 10 mM NaCl, and 10% glucose at pH 8.0. Localization drifts were corrected with n autocorrelative algorithm^63^. The drift-corrected coordinates were converted to be compatible to Vutara SRX software by an Excel macro. Cluster analysis and pair correlations were determined with the default modules in Vutara SRX software. DBSCAN algorithm was applied to determine the clusters which are within the search radius (ε) of 100 nm and consisting of at least 10 localizations. The apparent raft size was calculated by measuring the full width half max (FWHM) of the clusters.

### Sandwich ELISA assay

HEK293T cells were cultured in 96-well cell-culture plates. Each well was incubated with and without 100 μL treatments for 1 h, then washed with 100 μL PBS once and incubated with 100 μL PBS for 1 h. Supernatants were collected and analyzed for Aβ40 ELISA.

A 96-well plate was coated with 50 μL capture antibody (IBL #11088) at 5 μg/ml concentration in PBS and incubated overnight at 4°C. All of the rest incubations were performed at room temperature. The plate was washed with 200 μL PBS for three times, and 100 μL blocking buffer (PBS with 10%BSA and 0.05% TritonX-100) was added to each well and incubated for 1 h. Next, the blocking buffer was removed, and 50 μL of supernatant was added to each well and incubated for 1 h, followed by an addition of 50 μL primary antibody (Invitrogen™ #PA3-16760) at 1:10000 dilution in PBST buffer (PBS with 0.01% TritonX-100). After a 3 h incubation, the plate was washed with 200 μL PBST for 4 times and 100 μL HRP-linked goat anti-rabbit IgG secondary antibody (Invitrogen™ #31460) at 0.4 μg/ml concentration in PBST buffer was added for 1 h incubation in the dark. Then, the plate was washed with 200 μL PBST for 4 times. 80 μL Chromogen (Invitrogen™ #002023) was added and incubated in the dark for 30 min. Finally, 80 μL stop solution (Invitrogen™ #SS04) was applied to terminate the substrate development. Measurement of absorbance at 450 nm was performed on a microplate reader (Tecan Infinite 200 PRO) to determine relative Ab40 concentration.

### Membrane Fluidity Test

Change of membrane fluidity of HEK 293T cells was measured using the Membrane Fluidity kit (Abcam) following the manufacturer’s protocol. Briefly, ~10,000 cells were seed in 96 well plates and incubated with the drugs and the fluorescent lipid reagent containing pyrenedecanoic acid (2 mM) at the room temperature for 20-30 mins. with. Pyrenedecanoic acid exists as either a monomer or an excimer, the latter forms due to the change in the membrane fluidity. The formation of the excimers shifts the emission spectrum of the pyrene probe to the longer wavelength. The changes in spectrum emission were measured with a fluorescence microplate reader (Tecan Infinite 200 Pro). The ratio of monomer (EM 372 nm) to excimer (EM 470 nm) fluorescence was calculated to obtain a quantitative change of the membrane fluidity.

### *In vivo* and *in vitro* PLD Assay

*In vivo* PLD2 activity was measured in cultured HEK 293T cells by an enzyme-coupled product release assay using amplex red reagent as described previously^14^. Cells were seeded into 96-well plates (~5×10^4^ cells per well) and incubated at 37 °C overnight to reach confluency. The cells were starved with serum-free DMEM for a day and washed once with PBS (phosphate-buffered saline). The PLD reaction was initiated by adding 100 μL of reaction buffer (100 μM amplex red, 2 U/ml horseradish peroxidase (HRP), 0.2 U/ml choline oxidase, and 60 μM C8-PC, 50 mM HEPES, and 5 mM CaCl2, pH 8.0). The assay reaction was performed for 2-4 hour at 37 °C and the activity was kinetically measured with a fluorescence microplate reader (Tecan Infinite 200 Pro) at excitation and emission wavelengths of 530 nm and 585 nm, respectively. For *in vitro* assay, cabbage PLD was used instead of the live cells and the PLD reaction was initiated as described for the *in vivo* assay. The PLD2 activity was calculated by subtracting the background activity (reaction buffer with the drugs, but no cells). For the bar graphs, samples were normalized to the control activity at the 120 min time point.

### Statistical Analyses

Data calculations and graphs were performed in Prism 6 (GraphPad software) or Microsoft Excel. Experiments were done two-three times to ensure reproducibility. All Experimental samples were performed in random orders when to avoid any experimental bias. To ensure the reproducible effect of the sample sizes, super resolution imaging was carried out on multiple cells. Statistical significance was evaluated using ANOVA with post hoc Dunnett’s test, two-tailed t-tests, parametric or nonparametric wherever appropriate. Data are shown as the mean and the error bars with SD. Significance is indicated by *P ≤ 0.05, **P ≤ 0.01, ***P ≤ 0.001, and ****P ≤ 0.0001.

## CONTRIBUTIONS

ZY and HW performed viral entry assays, ZY and MAP performed imaging experiments, and ZY performed the Aβ experiments with the help of HW. ZY, MAP, and SBH designed the experiments and wrote the manuscript.

## Acknowledgements

We are grateful Mike Farzan for providing SARS2-PV.

## Supplemental Figures

**Fig. S1.**
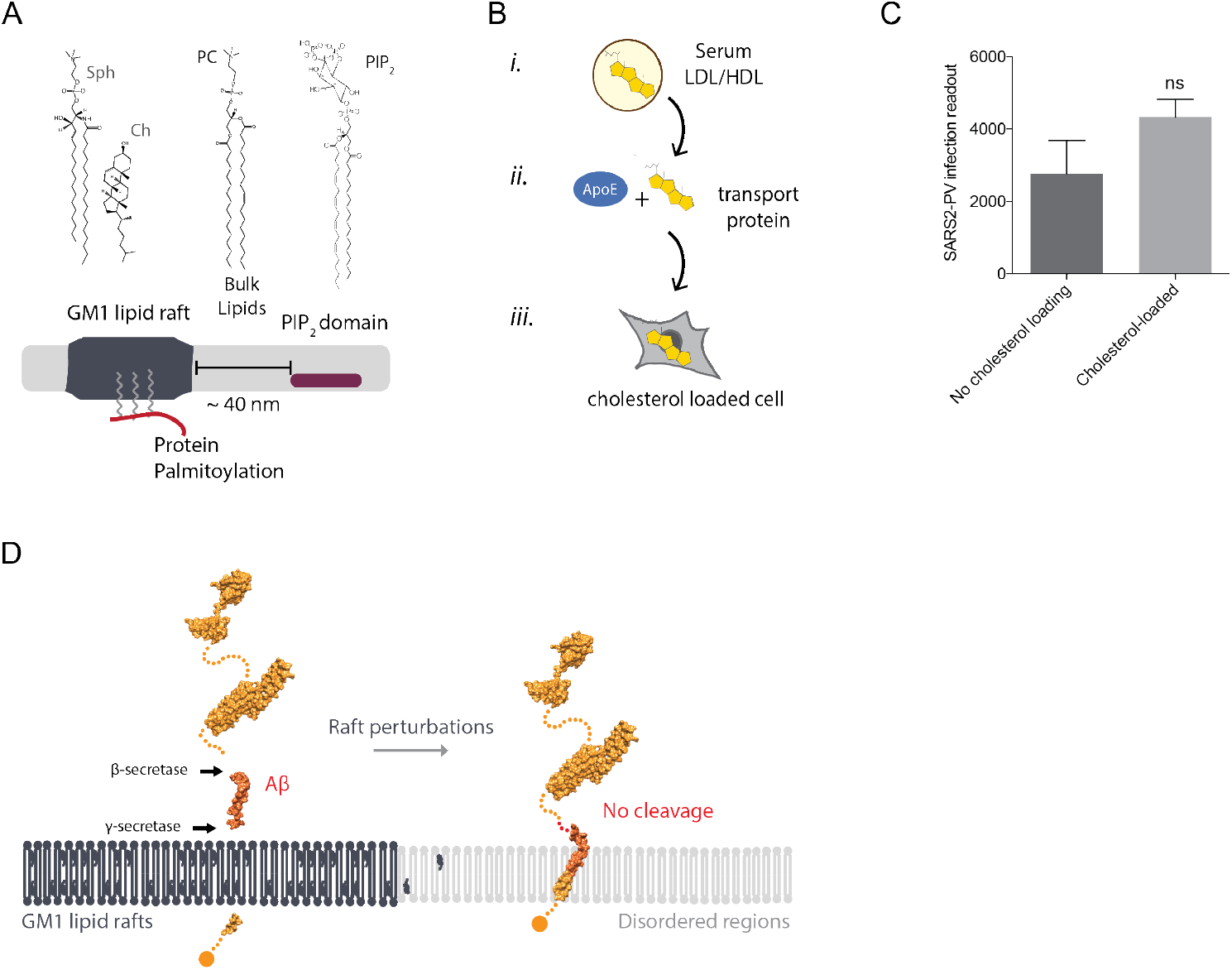
Membrane heterogeneity. **(A)** GM1 rafts are clusters of saturated known as liquid ordered (L_o_) and commonly reside separate from liquid disordered (L_d_) phases^62^. The ordered phase (L_o_) is generally enriched in sphingomyelin and cholesterol whereas the disordered (L_d_) phase consists of unsaturated lipids and includes polyunsaturated lipids like PA and PIP_2_ ^64^. **(B)** Cartoon diagram showing the experimental setup for loading cultured cells with cholesterol. ***i.***, Cholesterol (yellow shading) loaded into lipoprotein (e.g., low- and high-density lipoprotein (LDL and HDL respectively)) from blood serum. ***ii.***, Cholesterol free human apolipoprotein E (apoE, brown shading), a cholesterol transport protein, is exposed to cholesterol from blood serum and ***iii,*** ApoE transports cholesterol into of cells (grey shading). **(C)** SARS2-PV entry in ACE2 overexpressing HEK293T cells without and with cholesterol loading indicated by raw luciferase activity readout. Data are expressed as mean ± s.e.m., unpaired t-test, n=4. **(D)** Model of HCQ and anesthetics translocating APP from GM1 rafts to disordered regions through raft perturbation to reduce the synthesis of Aβ.

**Fig. S2.**
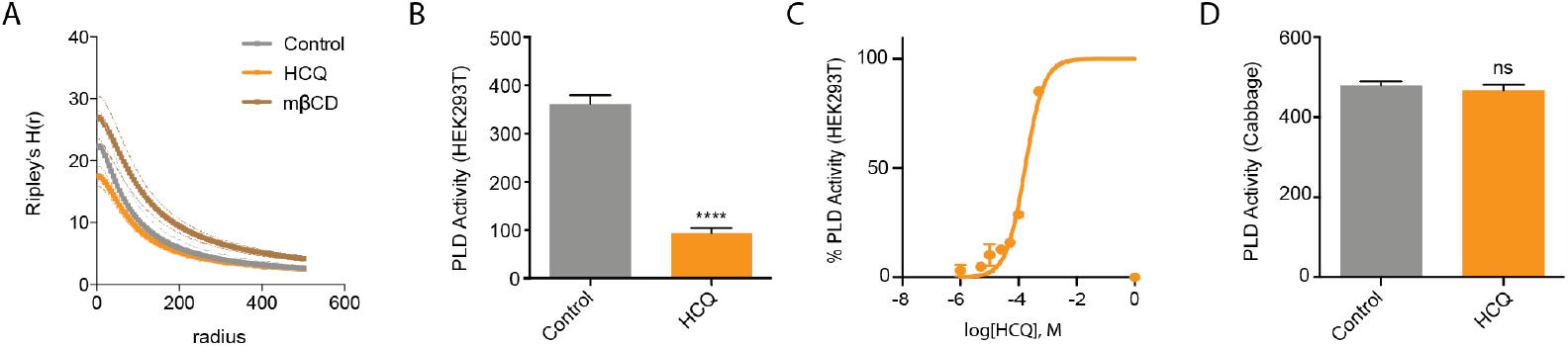
HCQ displacement of PLD2 from lipid rafts. **(A)** Ripley’s H-Function (H(r)) showing raft separation. **(B)** HCQ (50μM) decreased PLD activity in PLD assay. Data are expressed as mean ± s.e.m., ****P ≤ 0.0001, unpaired t test, n=6. **(C)** A dose response of HCQ’s inhibition to PLD activity in PLD assay, n=3. **(D)** Effect of HCQ(50μM) on PLD activity in cabbage PLD assay is not significant, unpaired t test, n=4-5.

**Fig. S3.**
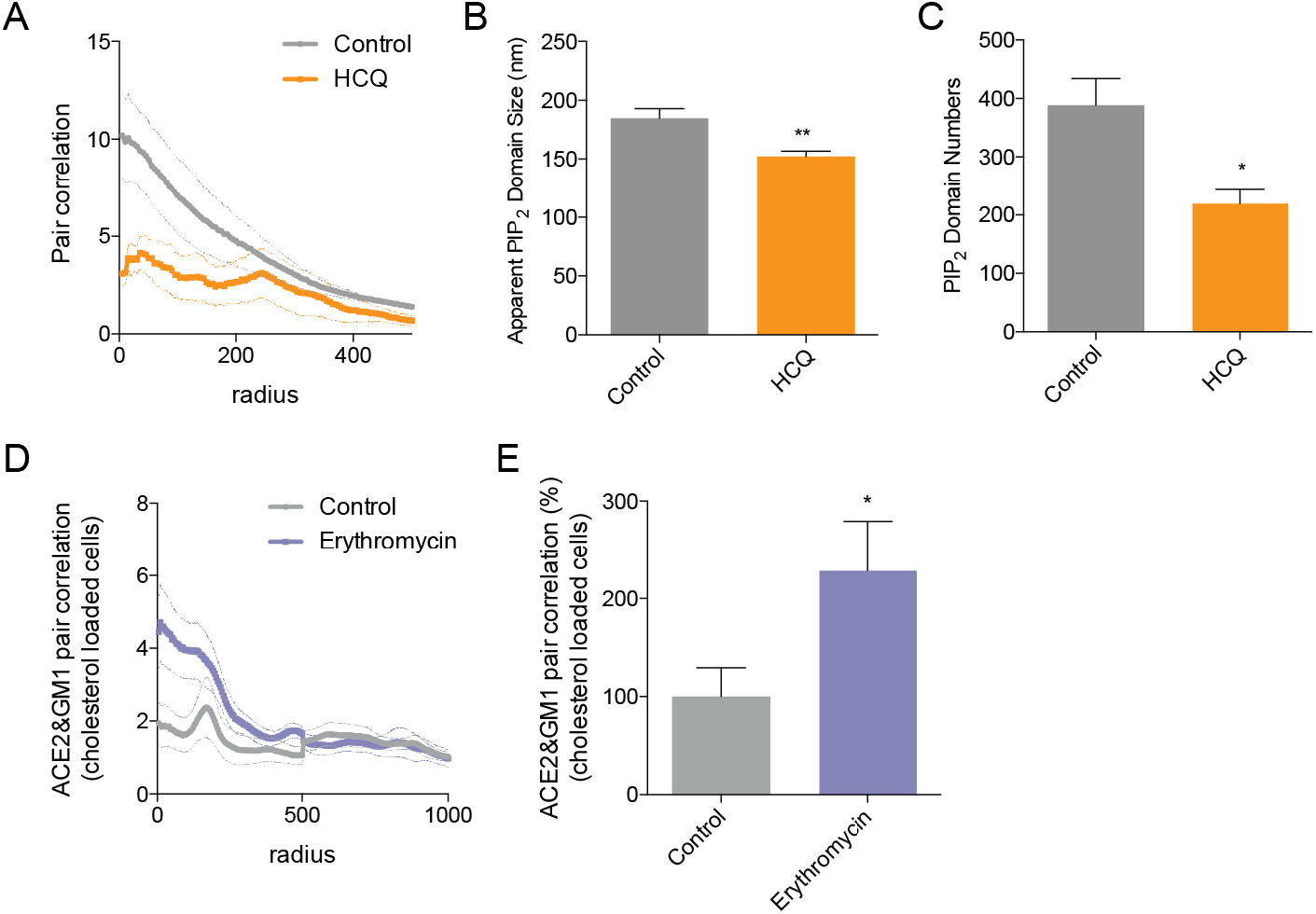
dSTORM of PIP_2_ domains. (**A**) Pair correlation analysis of dSTORM imaging (Fig. 3C). HCQ treatment decreased association of ACE2 and PIP_2_. **(B-C)** Bar graph of the apparent raft diameter analyzed by DBSCAN cluster analysis. HCQ decreases both raft diameter (B) and number (C) of PIP2 domains. Data are expressed as mean ± s.e.m., *P ≤ 0.05, **P ≤ 0.01, one-way ANOVA, n=5-6. **(D-E)** Pair correlation (D) and percent of pair correlation calculated at short distances (0-5 nm) (E) of dSTORM imaging. Erythromycin treatment decreased association of ACE2 with GM1 rafts. Data are expressed as mean ± s.e.m., *P ≤ 0.05, **P ≤ 0.01, unpaired t test, n=10.

